# Functionally annotating cysteine disulfides and metal binding sites in the plant kingdom using AlphaFold2 predicted structures

**DOI:** 10.1101/2022.10.17.512470

**Authors:** Patrick Willems, Jingjing Huang, Joris Messens, Frank Van Breusegem

## Abstract

Deep learning algorithms such as AlphaFold2 predict three-dimensional protein structure with high confidence. The recent release of more than 200 million structural models provides an unprecedented resource for functional protein annotation. Here, we used AlphaFold2 predicted structures of fifteen plant proteomes to functionally and evolutionary analyze cysteine residues in the plant kingdom. In addition to identification of metal ligands coordinated by cysteine residues, we systematically analyzed cysteine disulfides present in these structural predictions. Our analysis demonstrates most of these predicted disulfides are trustworthy due their high agreement (~96%) with those present in X-ray and NMR protein structures, their characteristic disulfide stereochemistry, the biased subcellular distribution of their proteins and a higher degree of oxidation of their respective cysteines as measured by proteomics. Adopting an evolutionary perspective, zinc binding sites are increasingly present at the expense of iron-sulfur clusters in plants. Interestingly, disulfide formation is increased in secreted proteins of land plants, likely promoting sequence evolution to adapt to changing environments encountered by plants. In summary, Alphafold2 predicted structural models are a rich source of information for studying the role of cysteines residues in proteins of interest and for protein redox biology in general.

## INTRODUCTION

Cysteine is a reactive sulfur-containing amino acid often residing in functional protein sites and metal binding sites. The cysteine content in proteomes is positively correlated with species complexity, ranging from 0.5% of all residues in prokaryotic proteins up to ~2% to 2.3% in higher eukaryotes [1–3]. In line with its functional importance, cysteine is one of the most conserved protein residues, especially in structurally buried cysteines buried cysteines and cysteines part of disulfides [3, 4]. Highly conserved cysteines are often part of CXXC motifs that facilitate enzymatic redox reactions [5] or binding of metal ligands [6, 7]. Metal ligands such as zinc (Zn^2+^) and iron-sulfur (Fe-S) clusters are mostly coordinated by cysteine and histidine residues [8, 9] and are often crucial to protein structure and function. Moreover, metal binding itself can be reversible and steer protein function dependent on the local redox homeostasis [10, 11]. Other dynamic redox switches of protein function are oxidative post-translational modifications resulting from the reaction of cysteine thiols with reactive oxygen/nitrogen/sulfur species [12].

Cysteine thiols also contribute to protein stability by formation of intramolecular disulfides. Within proteins, there are two categories of disulfides bonds [13]. The major one contains structural disulfides that are enzymatically catalyzed in plants via disulfide relay systems within the endoplasmic reticulum (ER), Golgi apparatus, the mitochondrial intermembrane space and, unique to plants, the thylakoid lumen of the chloroplast [14–16]. The second category entails functional disulfides including catalytic disulfides involved in thiol redox regulation, such as in thioredoxin domains, and allosteric disulfides that trigger conformational and possibly functional changes within a protein [17]. Next to intramolecular disulfides, cysteines can form intermolecular or so-called mixed disulfides between protein chains or with non-proteinaceous thiol compounds such as glutathione. Mixed disulfides typically occur during catalytic reduction reactions and redox relays or they can facilitate hydrogen peroxide dependent multimerization [18, 19] and even phase separation events in plants [20].

Despite significant advancements, proteome-wide identification of disulfide peptides with mass spectrometry remains challenging and would greatly benefit from more efficient enrichment methods [21]. Currently, systematic studies of disulfides in protein typically rely on available experimental structures [3, 22]. The release of AlphaFold2, however, has caused the sudden availability of highly accurate protein structures for entire proteomes [23, 24], recently releasing more than 200 million protein structures [25]. This presents now an unprecedented source to structurally annotate entire proteomes. Here, we leveraged AlphaFold2 predictions to identify disulfides and cysteine-dependent metal binding sites in fifteen proteomes of the plant kingdom.

## MATERIAL AND METHODS

### Protein sequences, phylogeny and orthology

Plant species diverging times were derived from TimeTree.org [26]. Small adaptations were the divergence of *Chlamydomonas* and *Ostreococcus* estimated at ~1,050 million years ago (MYA) [27] and *Physcomitrium patens* (moss) and *Marchantia polymorpha* (liverwort) at ~ 450 MYA [28]. All protein sequences used in this study were derived from UniProtKB reference proteomes (Table S1). Orthology relationships were determined using the PLAZA integrative orthology tool [29].

### Protein domains and subcellular localization

Protein domains from the InterPro database [30] were retrieved via the UniProtKB protein annotation. N-terminal sorting signals were predicted using the stand-alone version of TargetP 2.0 [31] using the default settings for plant proteins. For *Arabidopsis thaliana*, the SUBA database was used for subcellular protein annotation [32].

### Comparison to annotated disulfides and metal binding sites from PDB and UniProtKB

To benchmark disulfides from AlphaFold2 predicted structures, we compared to all available X-ray and NMR structures of PDB database (September 2022) for the fifteen plant species (Dataset S2). PDBrenum [33] was used to re-number residues in PDB structures according UniProtKB sequences (and thus AlphaFold2 residue numbering). Afterwards, SSBOND records were parsed from 2,602 PDB files. In addition, cysteines within PDB structures that were not part of a disulfide were kept track of in order to discern situations where AlphaFold2 did predict a disulfide. To compare metal ligand binding predictions, we retrieved cysteines that coordinate metal ligands (Feature Type: Sites, ‘Binding site’) in *Arabidopsis* as annotated by UniProtKB (‘Sites’ Feature Type: ‘Binding site’; Dataset S3).

### Computational processing of protein structures

Predicted protein structures were extracted from the online AlphaFold2 resource (https://alphafold.ebi.ac.uk/) and experimental structures from PDB (https://www.rcsb.org/). For computational processing of AlphaFold2 predicted structures we used the Biopython [34] PDB module to read in PDB structures and NumPy [35] to calculate geometric distances and dihedral angles in the Python3 programming language. The DSSP algorithm was used to assign secondary structure and relative solvent accessibility [36]. For pKa calculation, we used the PropKa3.0 algorithm [37]. The average depth of cysteine residues or its sulfur atoms was calculated using the MSMS algorithm [38]. All protein structure visualizations were created using ChimeraX (version 1.14). AlphaFold2 predicted and experimental X-ray or NMR structures were superimposed using the built-in matchMaker algorithm using default settings. For the ORF8 homodimer prediction, AlphaFold-Multimer (version 2.2.2) [39] was used using the full sequence database and using the highest ranked structural model. Metal ligands were predicted using the published metal ligand search algorithm (https://github.com/Elcock-Lab/Metalloproteome) [40].

### Data and source code availability

All the data generated here, together with the source code used for processing predicted protein structures is available at the GitHub repository https://github.com/willems533/PlantDisulfides.

## RESULTS

### The protein cysteine content increased during plant land colonization

To explore the relative cysteine content within plant proteins, we selected fifteen plant species from unicellular algae to higher land plants that are representative for plant evolution and all have annotated UniProtKB reference proteomes (Figure 1; Table S1). These include three green algae, two chlorophyte algae *Chlamydomonas reinhardtii* and *Ostreococcus tauri*, as well as the streptophyte algae *Chara braunii* that serves as an excellent model system for studying plant adaptation to land [41]. Land plant species include the bryophytes moss (*Physcomitrium patens*) and liverwort (*Marchantia polymorpha*), the vascular model system *Selaginella moellendorfii* and the basal angiosperm *Amborella trichopoda*. In addition, we included the widely studied model species *Arabidopsis thaliana* and *Populus tricochocarpa*, as well as economically important crops such as maize, rice, tomato and soybean. Of note, we also included eelgrass (*Zostera marina*), an angiosperm that re-adjusted to a marine habitat [42]. Taken together, all these plant species are representative for the plant kingdom, including plant evolutionary adaptations to terrestrial habitats and exposure to higher atmospheric oxygen concentrations (Figure 1).

**Figure 1.**
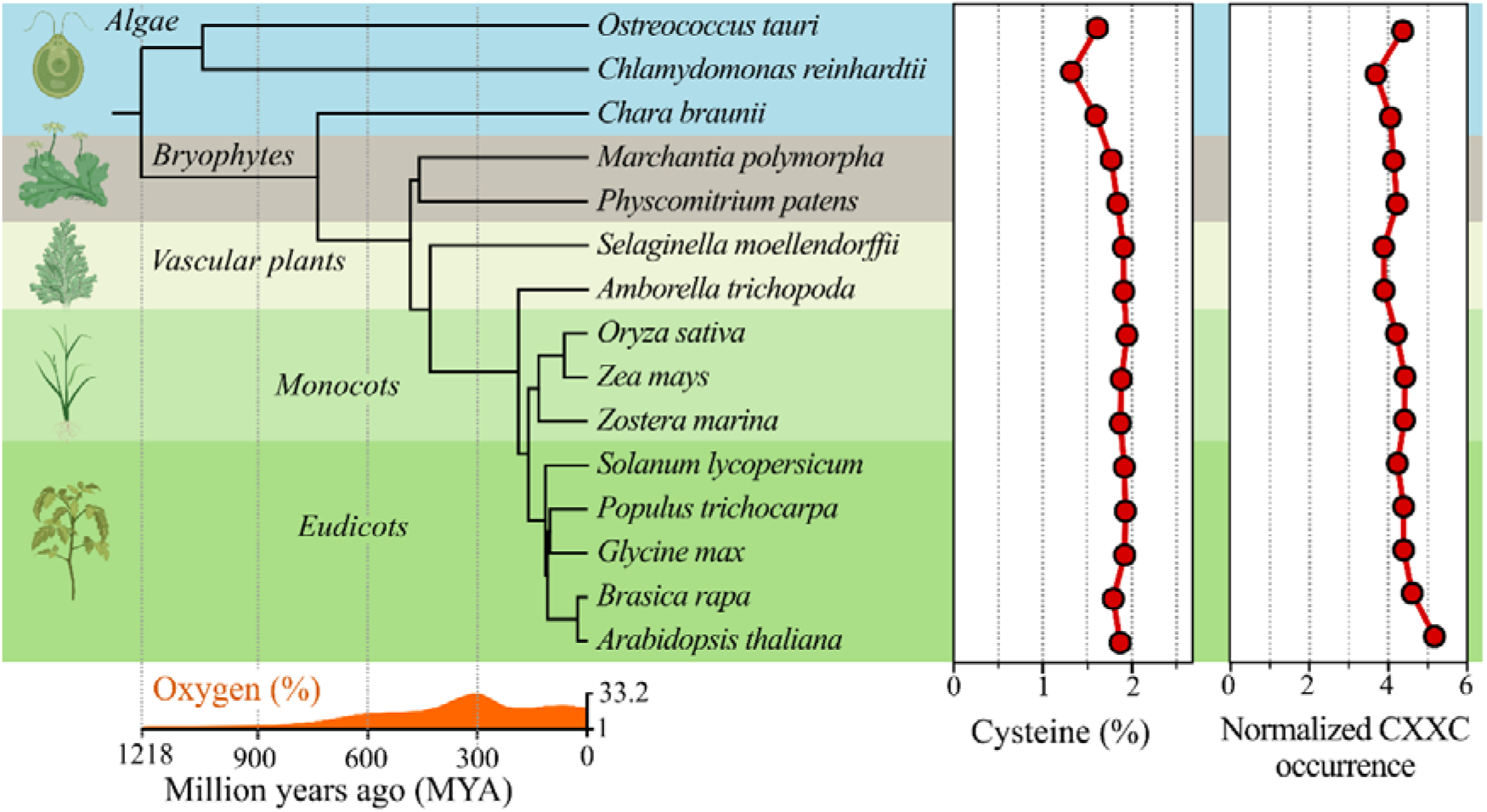
Cysteine and redox motif distribution in *Viridiplantae*. (*Left*) Timetree of fifteen plant species with divergence times (x-axis, million years ago [MYA]) and respective atmospheric oxygen concentrations (%) derived from TimeTree [26] with minor adaptations (see Methods). Plant graphics created with BioRender.com. (*Middle*) Percentage of cysteine residues in the UniProtKB reference proteome. (*Right*) Normalized occurrence of CXXC motifs. The number of motifs was divided by the number of cysteine residues in the proteome and multiplied by a hundred.

The percentage of cysteine residues shows a progressive increase from ~ 1.5% in algae towards ~ 1.9% to 2.0% in land plant proteomes over a period of approximately 1.2 billion years (Figure 1, Table S2). Counterintuitively, *Chlamydomonas* and *Chara* present a higher number of cysteines per protein due to the in general larger protein size with an average protein length of 650 to 740 amino acids in these species (Figure S1). Next to mere cysteine content, we also analyzed the occurrence of the CXXC redox motif relative to the number of cysteine residues within each proteome (Figure 1). Despite their lower cysteine content, algae present similar normalized numbers of CXXC motifs compared to land plants. In all plants, cysteine is preferably positioned within CXXC motifs compared to CC to CX_5_C motifs (Figure S2). Hence, a gradual increase of proteome cysteine content is evident throughout plant evolution, although, relative to its cysteine content, CXXC redox motifs were already prominent in algae species.

### Metal coordination by cysteine increases throughout plant evolution

Next, we investigated the structural context of cysteines within 430,374 AlphaFold2 predicted structures for the fifteen plant proteomes (Dataset S1). Note that for approximately 3% (15,351) of the UniProtKB reference proteins no AlphaFold2 structures are available (e.g. due to protein length constraints) or lack a cysteine residue (29,795 proteins, 6.27%). First, we identified cysteines residing in Zn^2+^ and Fe-S cluster binding sites in all the predicted structures by using a highly specific metal ligand searching algorithm [40]. Briefly, this algorithm iterates over possible metal binding sites in a structure and superimpositions each Fe-S and Zn^2+^ ligand, retaining the best scoring ligand in terms of root mean square deviation and steric clashes. Except for *Ostreococcus* (6.8%), the number of metal binding cysteines ranges between 4.5 to 6% in algae and bryophytes, while more than 6% in angiosperms – peaking at 9.6% in *Arabidopsis* (Figure 2). In all plant species the majority (> 90%) of metal binding cysteines coordinate Zn^2+^, while the other fraction participate in Fe-S clusters. Within the latter fraction, cysteines are more prevalent in algae and bryophytes, in contrast to Cys_2_His_2_ Zn^2+^ binding sites that are more present in higher land plants (Figure S3). Similarly, metal binding protein domains such as Cys_2_His_2_ zinc finger domains are more prevalent in land plants and, oppositely, lower proportions of Fe-S cluster containing domains, such as the [4Fe-4S] cluster in the radical SAM domain [43], are more abundant in green algae (Figure S4).

**Figure 2.**
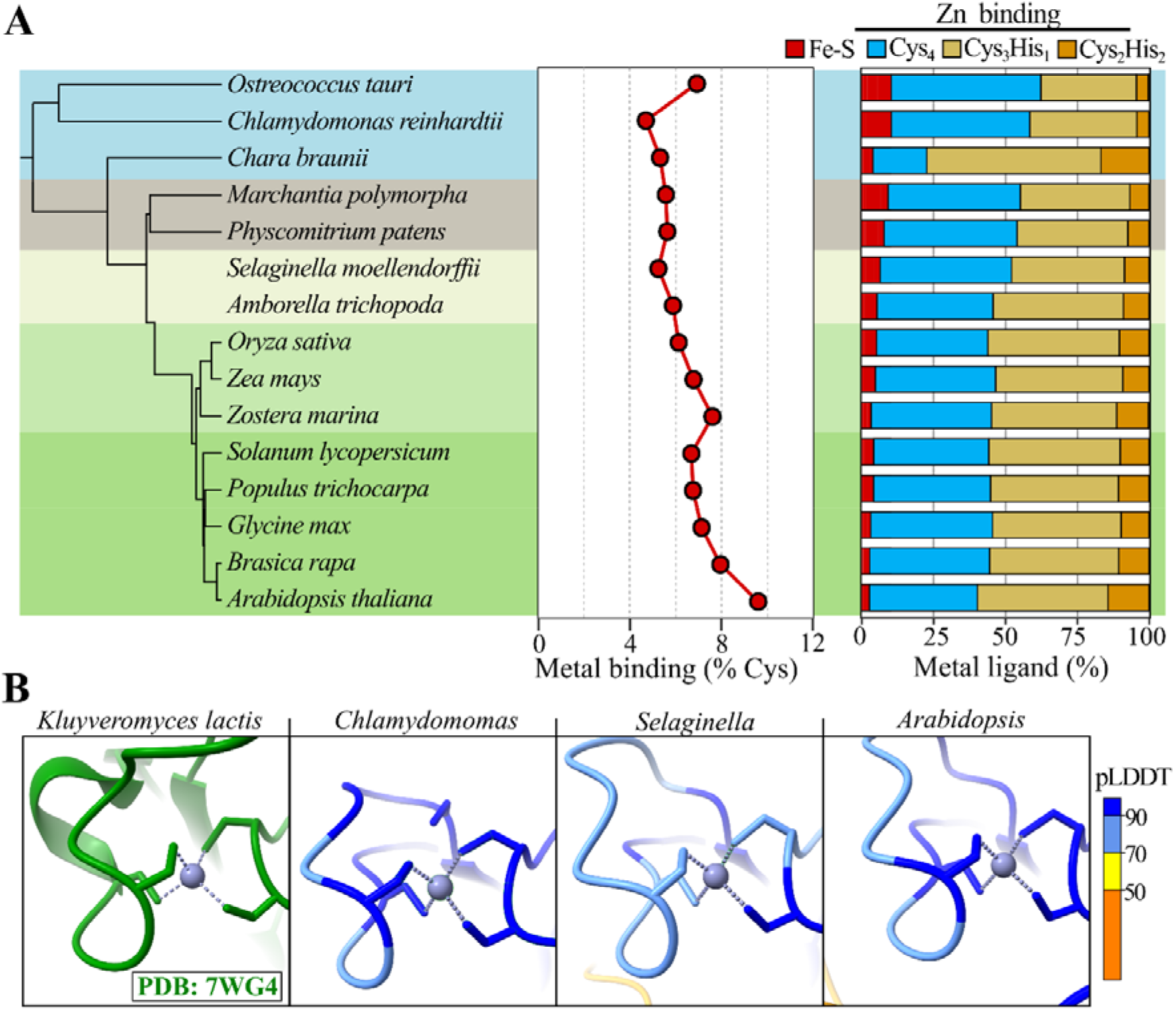
Metal coordinating cysteine motifs in plants. (**A**) (*Left*) Schematic representation indicating species relationships. (*Middle*) Percentage of cysteine residues participating in metal ligand coordination based on AlphaFold2 predicted protein structures using the method of [40]. (*Right*) Distribution of cysteines involved in different metal ligand binding sites. For an overview per metal ligand see Figure S3. (**B**) Cys_4_ Zn^2+^ binding site in *Kluyveromyces lactis* Arg-tRNA-protein transferase 1 (Ate1) (PDB 7WG4, green) [44] and AlphaFold2 predicted structures for *Chlamydomonas* (UniProtKB A0A2K3D0B7), *Selaginella* (D8RCG9), and *Arabidopsis* (Q9ZT48) orthologs. Zn^2+^ ligands were modeled by the metal ligand searching algorithm [40]. AlphaFold2 residues were colored according the per-residue confidence score (pLDDT) with > 90 being “highly confident” and < 70 being “low confident” predictions.

To showcase the potential relevance of these metal binding site predictions, we turned to a recently discovered metal ligand binding site in the arginyl-tRNA-protein transferase (ATE), which is conserved in eukaryotes. Whereas binding studies using recombinant *Saccharomyces cerevisiae* Ate1 indicate the presence of Fe-S clusters [45], the crystal structure of recombinant *Kluyveromyces lactis* Ate1 (Figure 2B – *Left*) identified instead a Cys_4_ Zn^2+^ binding site in the corresponding binding pocket [44]. In the AlphaFold2 predicted structures of both yeast Ate1 as well as their plant orthologs, we identified consistently a Cys_4_ Zn^2+^ binding site, like shown in Figure 2B for *Chlamydomonas, Selaginella* and *Arabidopsis*, and thus alike those reported for *Kluyveromyces lactis*. As such, AlphaFold2 predicted structures of plant proteins can be powerful hypothesis generators, predicting here a recently discovered metal binding site that might be of functional relevance to plant ATE enzymes functioning in N-degron pathways and oxygen sensing [46, 47].

### A characteristic disulfide chemistry is present in AlphaFold2 predicted structures

Disulfide bonds are on average 2.05 Å in length and within protein structures a threshold of 2.5 Å has been used for systematic structural analysis of disulfides [22] or algorithms such as the *p*K_a_ predictor PROPKA [48]. Applying this distance constraint of 2.5 Å, 195,439 intramolecular disulfides were retrieved for AlphaFold2 predicted protein structures of the fifteen plant species (Dataset S1). Both the disulfide length (median 2.03 Å, Figure 4A) and *Cα* atom distance (median 5.52 Å, Figure S5A) matched with anticipated disulfide length of ~ 2.05 Å and 5.6 Å, respectively [49, 50]. Moreover, the five dihedral angles (χ1, χ2, χ3, χ2′, and χ1) defined by the atoms underlying the disulfide bond (Figure 3B) agreed with those derived from experimental data (Figure S5B-D). For instance, the χ3 torsion angle (formed by C_β_–Sγ–Sγ–C_β_ bonds) peaked at approximately −85° and +100° (Figure 3C), corresponding to left- and right-handed disulfides, respectively [50–52]. Taken together, this precise disulfide stereochemistry highlights the accuracy of cysteine sidechain predictions by AlphaFold2, which was also demonstrated by the correct orientation of up to four cysteine sidechains to form metal binding pockets [40].

**Figure 3.**
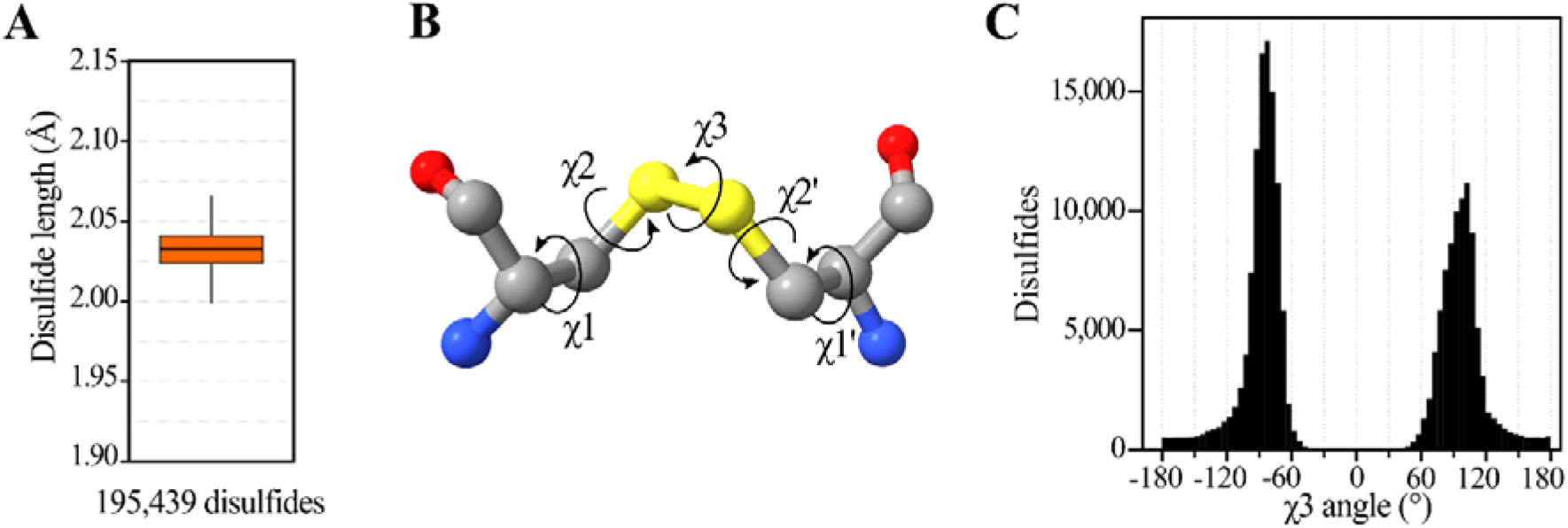
Stereochemistry of disulfides in AlphaFold2 predicted structures. **(A)** Disulfide bond (sulfur-sulfur atom) distance for 195,439 disulfides in AlphaFold2 predicted structures **(B)** Ball-and-stick model of a disulfide, indicating the five dihedral angles of a disulfide: χ1, χ2, χ3, χ2′, and χ1′. Carbon, nitrogen, oxygen and sulfur atoms were colored in grey, blue, red and yellow, respectively. **(C)** Histograms of χ3 dihedral angle (x-axis, binned per 5°) of disulfides in AlphaFold2 predicted structures. For other dihedral angels see Figure S5.

**Figure 4.**
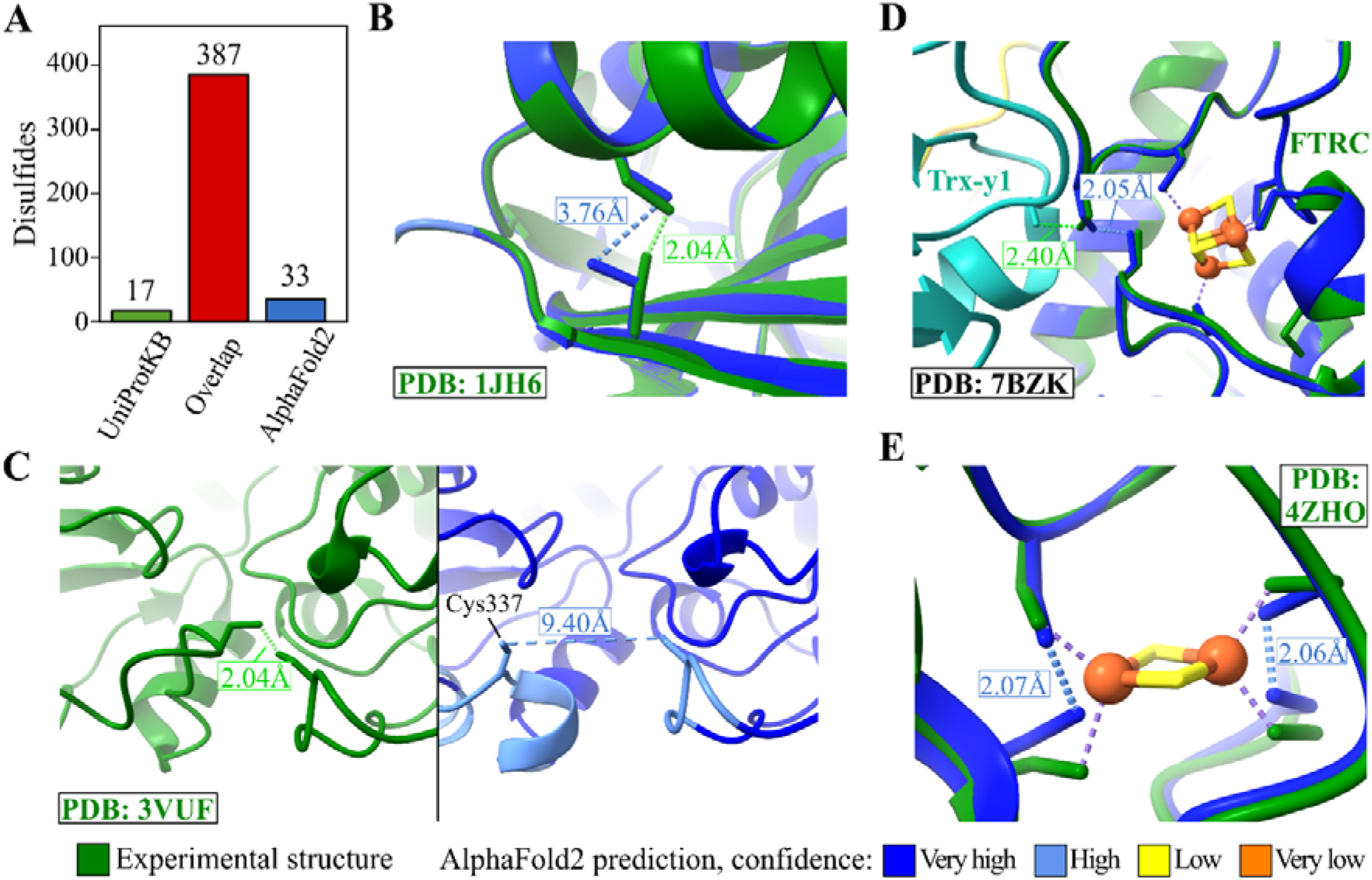
Comparison of AlphaFold2 predicted protein disulfides with experimentally determined disulfides. (**A**) Overlap of AlphaFold2 predicted disulfides and disulfides (‘SSBOND’) from experimental protein structures from PDB available for the fifteen plant species. (**B-E**) Comparisons of AlphaFold2 predicted and experimental protein structures with non-matching disulfides. Iron-sulfur clusters were determined only experimentally (E) or also predicted by the metal ligand searching algorithm in the AlphaFold2 predicted structure (D). AlphaFold2 predicted proteins were colored by each residue’s pLDDT score, ranging from dark blue (pLDDT > 90, “very high confidence”), light blue (pLDDT > 70, “confident”) and low and orange indicting a “low and very confidence” prediction, respectively.

### AlphaFold2 predicted disulfides are consistent with X-ray and NMR protein structures

To compare disulfides in AlphaFold2 predicted structures with experimental determined protein disulfides, we parsed 429 unique disulfides (‘SSBOND’ records) in 171 plant proteins using 2,602 available experimental structures deposited in the Protein Data Bank (PDB, https://www.rcsb.org/; Dataset S2). Notably, for 27 out of these 429 disulfides (6.3%), the pairing cysteines were also found as a free cysteine. Closer inspection learned that these cases mostly relate to small structural differences within different conformations of a protein (complex) or crystallization of oxidized and (semi-)reduced forms of a protein. Of the other cysteine pairs exclusively identified as disulfide, 387 (95.8%) were correctly predicted by AlphaFold2 (Figure 4A). The 17 experimental disulfides that were not present in AlphaFold2 predicted structures included eight cases with small deviations where Cys were ~ 3 to 5 Å apart, for instance due to a slightly different orientation of a single cysteine sidechain (Figure 4B). However, we observed also six cases with different backbone predictions, often coinciding with lower pLDDT confidence scores. For instance, Cys337 of rice STARCH SYNTHASE1 forms a disulfide with Cys529 but was 9.4 Å apart due to a different backbone prediction by AlphaFold2 that coincided with relatively lower confidence pLDDT scores (pLDDT < 90; Figure 3C). The remaining three wrongly predicted disulfides stemmed from lower resolution NMR structures. Note that when only considering disulfides from experimental protein structures not included in the AlphaFold2 training set (released after April 2018), 131 out of 137 disulfides (95.6%) were correctly assigned.

Next, we inspected the 33 disulfides predicted by AlphaFold2 lacking experimental validation in the experimental structures (Dataset S2). Similar to the above cases, sixteen cysteine pairs are within ~ 3 to 5 Å distance, and three more distantly spaced cysteines were due to deviating backbone predictions. In the case of FERREDOXIN-THIOREDOXIN REDUCTASE C (FTRC), the Cys87-Cys117 disulfide was absent in experimental structures of FTRC in complex with thioredoxins (Trxs), where Cys87 formed mixed-disulfides with Trx-*y1* (and Trx□*f2*, Trx□*m2*) (Figure 4D) [53]. However, prior to reaction with Trx, the nucleophilic Cys87 forms an intramolecular disulfide with Cys117 [54], thus consolidating the AlphaFold2 prediction. While the Fe-S cluster of FTRC was correctly superimposed in the AlphaFold2 predicted structure (Figure 4D), the [2Fe-2S] clusters in *Arabidopsis* and maize ferredoxin were not identified by the metal ligand search algorithm. Instead, AlphaFold2 formed two pair of disulfides between metal coordinating cysteines (Figure 4E). AlphaFold2 was reported before to occasionally forms disulfide bonds in metal binding sites due to the cysteine proximity [40]. In this regard, we did identified 960 cysteines that were within a 2.5 Å distance, but that also identified to coordinate a metal ligand, and hence as such categorized (Dataset S1). To further determine whether disulfides might actually be part of non-predicted metal binding sites, we compared them with the 2995 cysteines in *Arabidopsis* proteins annotated to bind Zn^2+^ or Fe-S clusters in UniProtKB. From these, 2669 cysteines (89.1%) were correctly assigned in our study and 55 (1.8%) not predicted as metal binding were classified as disulfides (including abovementioned ferredoxins) (Dataset S3). Hence, cysteines part of metal binding sites are adequately assigned in this study and hardly gave rise to incorrect disulfides in our analysis.

### Disulfide are overrepresented in secretory pathway proteins

Next, we assessed the amount of disulfides in different subcellular compartments. Therefore, we first categorized the subcellular location of proteins in all plant species based on the presence of N-terminal sorting signals predicted by TargetP 2.0 [31], scoring the presence of a secretory signal peptide (SP) and mitochondrial, chloroplast and thylakoid lumenal transit peptides (mTP, cTP and luTP, respectively). The prediction of N-terminal sorting signals ranged between 6.7% (*Chara*) up to 25.5% (*Arabidopsis*) per plant proteome (Figure S6). Next, we assessed the proportion of cysteine residues that form disulfides according the predicted N-terminal sorting signals. For proteins lacking N-terminal sorting signals, 6.20% of the cysteines form disulfides, while merely 3.99% and 3.01% in proteins containing cTP and mTP, respectively. Conversely, 57.8% of cysteines in proteins targeted to the secretory pathway form disulfides (with a predicted secretory SP), and 26.1% of those proteins targeted to the thylakoid lumen (Figure 5A – *Top*). These numbers are in line with the subcellular locations for oxidative folding of plant proteins [14, 15]. Disulfides have been identified in at least 30 thylakoid lumenal proteins [55], including VIOLAXANTHIN DE-EPOXIDASE (VDE) that contains six functionally important disulfide bridges [56], all of them present in the AlphaFold2 predicted with structure, with five of them in the N-terminal cysteine-rich domain (Figure 5C). Thus far, only a single disulfide, Cys231-Cys362, was identified in an experimental structure of the *Arabidopsis* VDE lipocalin protein domain (Figure 5C) [57]. In addition to the 30 reported disulfide-containing lumenal proteins [55], we identified thylakoid lumenal 17.9 kDa and protein nuclear-encoded photosystem II subunit T to contain disulfides in multiple plant species (Figure S7). While predicted sorting signals are a useful proxy for subcellular protein annotation of relatively less annotated plant species, we next relied on a more refined subcellular protein annotation for Arabidopsis. Based on the SUBA consensus location [32], 72.1% of cysteines residues of extracellular proteins form disulfides (Figure 5A – *Bottom*), approximately 20% in the vacuole, plasma membrane, and Golgi apparatus, and 15% of cysteines in proteins residing in the endoplasmic reticulum (ER). Conversely, proteins in the cytosol, mictochondrion, plastid, peroxisome and nucleus have a lower disulfide content (~2% to 4.5% of cysteines), which is in line with redox-sensitive biosensors based observations that demonstrate these compartments as reducing environments in *Arabidopsis*, unlike the oxidative ER [58].

**Figure 5.**
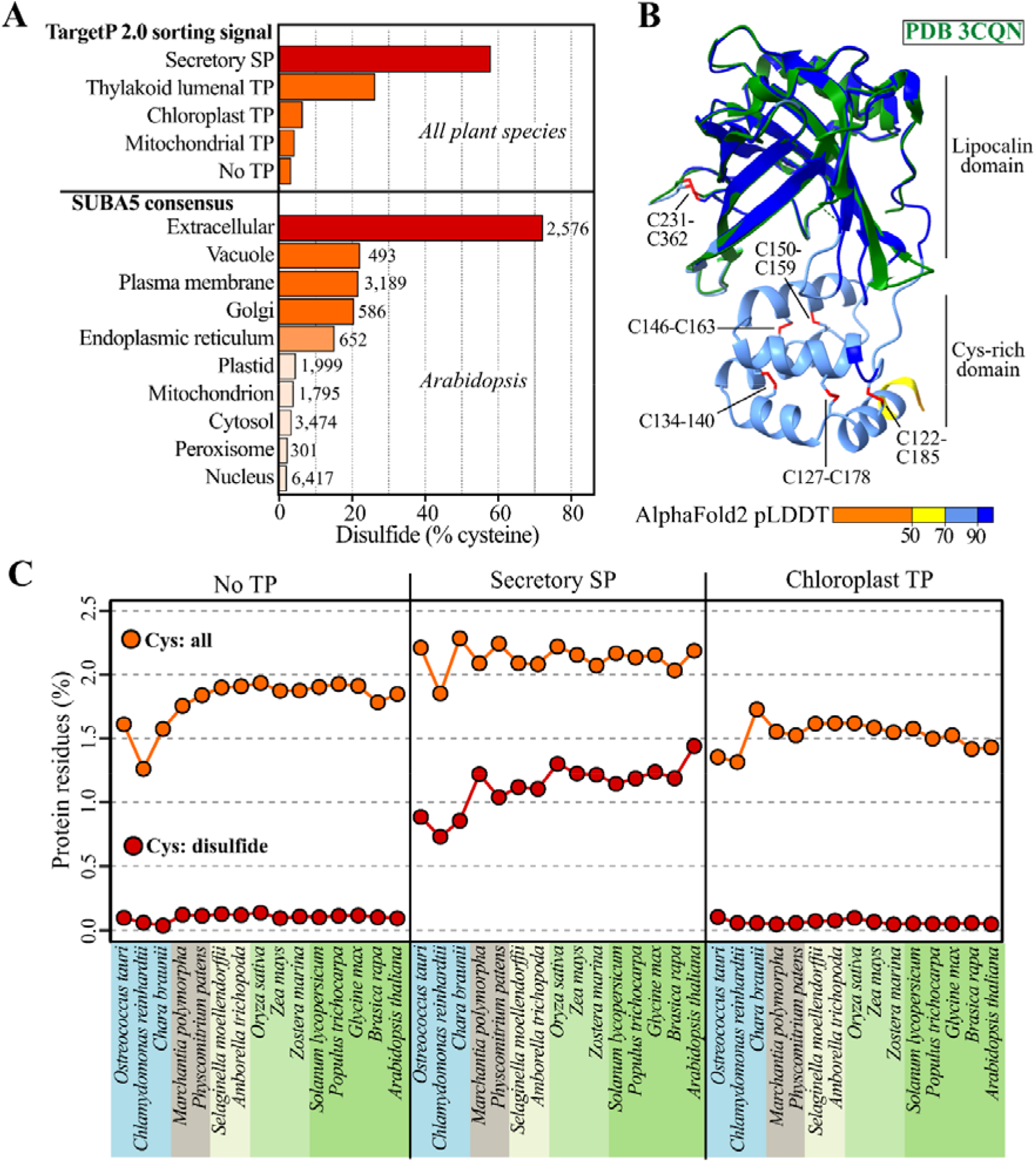
Cysteine and disulfides are (increasingly) abundant in secreted plant proteins. (**A**) Proportion of cysteine residues forming disulfides for proteins according (*i*) N-terminal sorting signal predictions by TargetP 2.0 for all plant species (*Top*) [31] and (*ii*) SUBA4 consensus locations for *Arabidopsis* proteins (with total protein numbers between parentheses) (*Bottom*). (**B**) Comparison of AlphaFold2 predicted and experimental protein structure (PDB: 3CQN; [57]) for VIOLAXANTHIN DE-EPOXIDASE (VDE; UniProtKB Q39249). Disulfides were indicated in red. AlphaFold2 predicted proteins were colored by each residue’s pLDDT score, ranging from dark blue (pLDDT > 90, “very high confidence”), light blue (pLDDT > 70, “confident”) and low and orange indicting a “low and very confidence” prediction, respectively. (**C**) Proportion of cysteine residues (orange) and cysteine residues forming a disulfide in AlphaFold2 predicted structures (red) in plant proteins according N-terminal sorting signal predictions. For all compartments see Figure S8. Abbreviations: SP, signal peptide; TP, transit peptide.

Given the strong subcellular preference of disulfides, we next assessed the proportion of all cysteine residues and those that form disulfides in a subcellular context. While proteins lacking sorting signals show increased cysteine proportions, secreted proteins have a cysteine content above 2% in all species except in *Chara* (Figure 5C). For other compartments, like the chloroplast, there appears a lower cysteine content of ~1.3% in the green algae *Chlamydomonas* and *Ostreoccus* compared to ~1.5% in the streptophyte algae *Chara* and land plants. Thus, cysteine content in secreted proteins does not correlate with complexity of plant species. In contrast, other subcellular compartments display a more variable cysteine content. While for proteins lacking a N-terminal sorting signal there is a increased cysteine content stagnating at ~1.9% in land plants (Figure 1), it only reaches up to ~1.6% in mitochondrial proteins (Figure 5C). Within the land plants, there is however an increased tendency towards disulfide formation in secreted proteins. A trend which is not apparent in proteins residing in other subcellular compartments (Figure 5C; Figure S8). These disulfides in land plant proteins are often involved in intercellular communication, cell wall processes or defense responses that for instance contain disulfide-containing domains such as pectinesterase inhibitor domains, secretory peroxidase domains or defensin-like proteins (Figure S9).

### Cysteines in disulfides and metal binding sites show a higher degree of oxidation

Two recent redox proteomics studies could quantify the percentage of cysteine oxidation in *Arabidopsis* cells via differential labeling of cysteines before and after a general thiol reduction step (e.g. DTT/TCEP), thereby discriminatively labeling both natively reduced thiols and reversibly oxidized thiols, respectively. In addition to disulfides, the reversibly oxidized thiols include various oxidative post-translation modifications such as *S*-sulfenylation, *S*-nitrosylation, persulfidation, and others. In the first study, iodoTMT isobaric labels were used to quantify rapid mitochondrial thiol reduction events reflective of seed imbibition [59]. In a second study, isotopic-labeled N-ethylmaleimide was used to quantify thiol oxidation changes in H2O2 treated leaves [60]. Next, we plotted the percentage of oxidation for cysteines categorized in our study to be either unbound, to be part of disulfides bonds or involved in metal ligand binding based on AlphaFold2 predicted structures. In isolated mitochondria, disulfide forming cysteines display on average 75% of oxidation, while 52 % oxidized in leaves (Figure 6). Hence, as anticipated, cysteines part of predicted intramolecular disulfides have a higher degree of oxidation. In case of metal binding cysteines, increased oxidation is only noted in the mitochondrial study (~ 45%) [59]. This apparent discrepancy is probably due to the acidic protein extraction buffer used for the leaves [60], which destabilizes metal binding sites and hence renders cysteines susceptible for alkylation prior to the thiol reduction step. Notably, cysteines categorized in our analysis to be unbound yet that appear to be highly oxidized (e.g. > 50%) might represent cysteines forming disulfides, binding metal ligands (or other cofactors), or simply be very susceptible to post-translational modifications. For example, Cys165 of MITOCHONDRIAL FERREDOXIN1 (MFDX1) was 72.5% oxidized and not predicted to be part of a disulfide or metal binding site based on AlphaFold2 predicted structures (Figure 6). However, this cysteine is known to bind a [2Fe-2S] cluster [61] and is reminiscent of the [2Fe-2S] cluster in chloroplast ferredoxins (Figure 4E), though cysteine pairs are now ~ 3.3 Å apart. Taken together, quantifying the degree of cysteine oxidation can offer complementary evidence to functionally annotate cysteines in combination with AlphaFold2 predicted structures.

**Figure 6.**
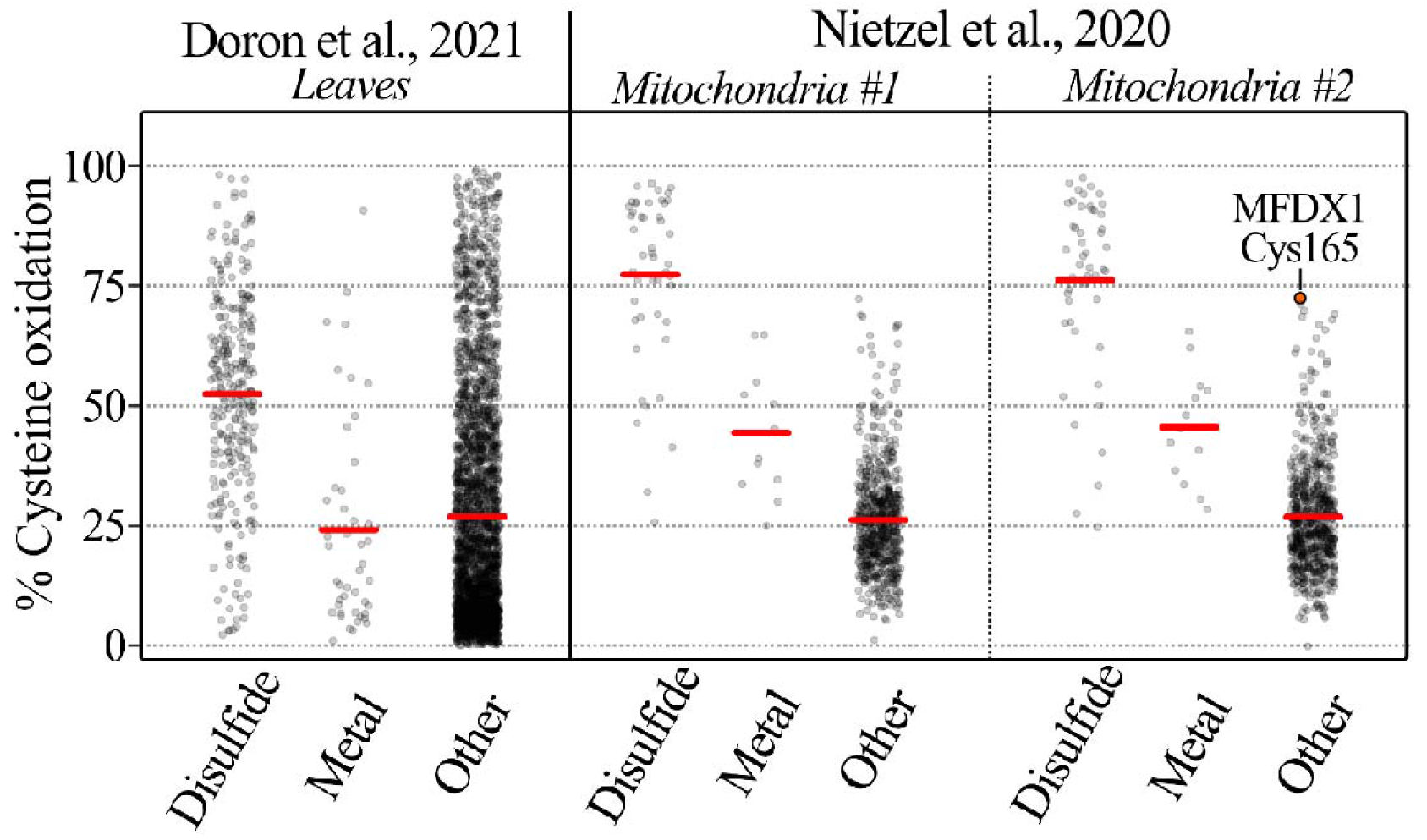
Metal binding and disulfide formation result in elevated cysteine oxidation measured by proteomics. The percentage of cysteine oxidation (y-axis) was plotted according cysteines annotated in this study as free cysteine or part of a disulfide or metal-binding sites in this study based on AlphaFold2 predicted structures (x-axis). The degree of cysteine oxidation (%) was derived from the supplementary data of two independent studies in *Arabidopsis* [59, 60]. Only peptides containing a single cysteine and quantified in at least two out three replicates of the control sample were considered. For the study of Nietzel et al., #1 reflects the control sample for the 2-oxoglutarate treatment and #2 the control sample for the citrate treatment. Abbreviations: MFDX1, MITOCHONDRIAL FERREDOXIN1.

## DISCUSSION

In this work we took advantage of the recent release of more than a million AlphaFold2 predicted protein structures [25] to structurally annotate all cysteines in fifteen plant proteomes. For the model land plant *Arabidopsis*, approximately 25% (50,312 / 208,029) of all cysteines were predicted to either form a disulfide (14.8% Cys) or bind metal ligands (9.6% Cys). All our analyses suggest that AlphaFold2 predicted disulfides are highly accurate. This is supported at multiple levels: a strong agreement (~96%) with disulfides previously identified in X-ray and NMR protein structures, a characteristic disulfide stereochemistry, a higher presence in oxidative subcellular compartments and a high percentage of cysteine oxidation as measured by proteomics.

The proportion of cysteines in a proteome positively correlates with the complexity of the species [1, 2]. Here, focusing on plants, a similar trend was observed with green algae (~ 1.5%), mosses (~ 1.7%) and higher land plants (~ 1.9%). However, this trend does not apply to the secreted proteins, which maintain a higher and relatively stable cysteine content of ~ 2 to 2.4%. In addition to a higher cysteine content, AlphaFold2 predicted structures of proteins routed via the secretory pathways are rich in disulfides, in line with experimental observations [22]. These disulfides are critical to stabilize the protein structure in the oxidative extracellular milieu [62]. Moreover, the increased structural stability offered by disulfides accelerates sequence evolution of membrane and extracellular proteins [63] that are evolving faster than cytosolic proteins due to a stronger selective pressure for adaptation to changing environments [64]. In this perspective, the observed increase in disulfide formation in land plants likely positively correlates with sequence evolution of secreted proteins and thereby facilitated adaptations to new terrestrial habitats, such as intercellular communication, cell wall biogenesis and pathogen defenses.

Using an established algorithm to search for metal ligands in AlphaFold2 predicted structures [40], we identified ~ 37,000 metalloproteins in fifteen selected plant species. This revealed that the proportion of cysteine to coordinate Zn^2+^ is elevated in land plant proteins, at the expense of Fe-S clusters that are more abundant in green algae. This phenomenon of increased Zn^2+^ binding has been observed before in eukaryotes compared to prokaryotes and has been linked to rising atmospheric oxygen concentrations [65, 66]. More concretely, oxygenic photosynthesis and consequently increased oxygen levels led to a decrease of soluble Fe^2+^ levels in oceans, opposed to transition metals such as Cu and Zn^2+^ that are estimated to have increased in concentration ~ 800 million years ago [67]. This is also reflected in the plant proteomes, with relatively more Fe-S clusters present in the chlorophyte algae *Ostreococcus* and *Chlamydomonas* that have been estimated to diverge from streptophyte algae and plants roughly around 725 to 1200 million years ago [68]. The freshwater streptophyte algae *Chara* presented an unusual case, with unusually high proportions of Zn^2+^ binding sites due to the presence of reverse transcriptases and nucleases that are not typically found in algae or land plants (Figure S4) [41]. While genomes and proteomes of other sequenced streptophyte algae would have been interesting to include in our analysis, e.g. *Mesostigma viride* and *Chlorokybus atmophyticus* [69], we solely retrieved a UniProtKB reference proteome (and AlphaFold2 predictions) for *Chara braunii*.

A current limitation of AlphaFold2 predictions is that they are monomeric and lack potential cofactors, interactors, or other relevant biomolecules that can adjust protein conformation *in cellulo*. However, structural predictions of multimeric protein complexes are becoming possible now with algorithms such as AlphaFold-Multimer [39], ColabFold [70], RoseTTAFold [71] or AF2Complex [72]. Intrigued by the fact that already intermolecular disulfides have been proposed based on of AlphaFold2 predicted structures [72, 73], we used AlphaFold-Multimer to predict the structure of the SARS-CoV-2 ORF8 homodimer that contains an intermolecular Cys20-Cys20 disulfide (Figure S10A) [74]. Despite being described as a very challenging protein [23], the three intramolecular disulfides in ORF8 chains were correctly predicted and Cys20 of both chains were 3.329 Å apart in the AlphaFold-Multimer predicted structure (Figure S10A). Hence, this suggests that structural predictions can be informative to propose possible intermolecular disulfide linkages between proteins. Structural predictions can also be informative to predict or gain mechanistic insights on cofactor or ligand binding [40, 75, 76]. For instance, next to metal ligand binding, algorithms such as AlphaFill provide structural models of cysteines forming covalent bonds with cofactors such as c-type heme (e.g. CYTOCHROME C-1; Figure S10B) or MoO_2_-molybdopterin in NITRATE REDUCTASE2 (Figure S10C). Another important consideration is that AlphaFold2 structures are static, while protein conformation and its disulfides can be dynamic and interchangeable (Hogg, 2020). This is as also evidenced by the absence or presence of disulfides in different deposited PDB structures of the same protein (Dataset S2). Moreover, such disulfide alterations, and oxidative modifications, are especially anticipated in conditions perturbing redox homeostasis, such as environmental stresses faced by plants. Despite these considerations, these available structural predictions are of particular interest to protein redox biology.

In summary, AlphaFold2 predicted structures provide a compelling resource to identify cysteines forming disulfides and metal binding sites in proteins or whole proteomes of interest. This structural perspective can serve as an excellent starting point in hypothesis formulation in protein redox biology and future developments are bound to offer us increasingly accurate structural perspectives on cellular redox regulation.

## ACKNOWLEDGEMENTS

The computational resources (Stevin Supercomputer Infrastructure) and services used in this work to run AlphaFold-Multimer were provided by the VSC (Flemish Supercomputer Center), funded by Ghent University, FWO and the Flemish Government – department EWI.

## SUPPLEMENTAL DATASETS AND CODE AVAILABILITY

Supplemental Datasets S1-3 and the source code used for this work is available at the GitHub repository https://github.com/willems533/PlantDisulfides. Separate datasheets for the functional cysteine annotation per plant species are available.

## SUPPLEMENTAL TABLES

**Table S1.**
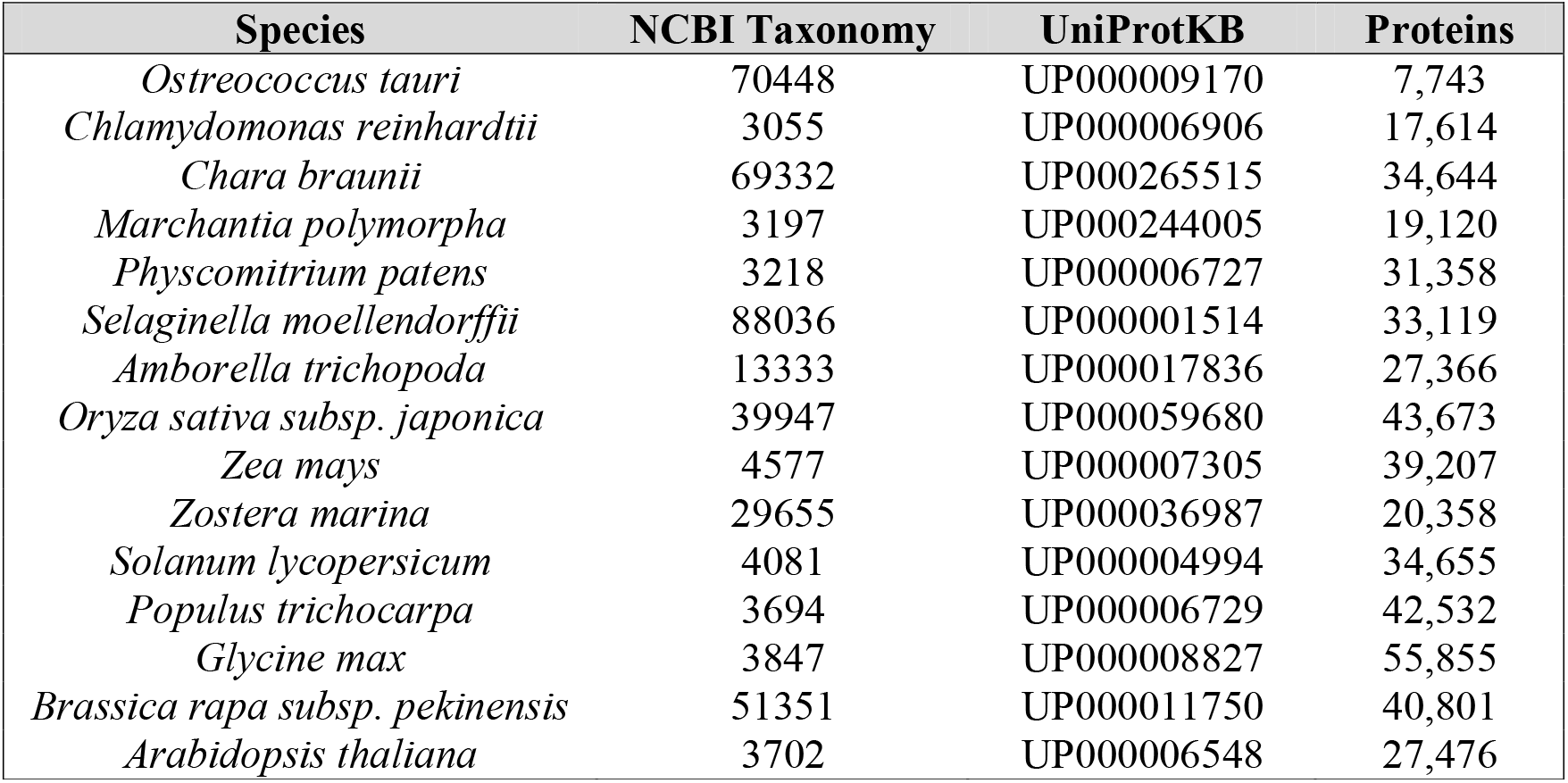
Fifteen plant species used in this study. The NCBI taxonomy identifier, UniProtKB reference proteome identifier and total of number of proteins per species. Proteomes were downloaded in September 2022.

| Species | NCBI Taxonomy | UniProtKB | Proteins |
| --- | --- | --- | --- |
| *Ostreococcus tauri* | 70448 | UP000009170 | 7,743 |
| *Chlamydomonas reinhardtii* | 3055 | UP000006906 | 17,614 |
| *Chara braunii* | 69332 | UP000265515 | 34,644 |
| *Marchantia polymorpha* | 3197 | UP000244005 | 19,120 |
| *Physcomitrium patens* | 3218 | UP000006727 | 31,358 |
| *Selaginella moellendorffii* | 88036 | UP000001514 | 33,119 |
| *Amborella trichopoda* | 13333 | UP000017836 | 27,366 |
| *Oryza sativa subsp. japonica* | 39947 | UP000059680 | 43,673 |
| *Zea mays* | 4577 | UP000007305 | 39,207 |
| *Zostera marina* | 29655 | UP000036987 | 20,358 |
| *Solanum lycopersicum* | 4081 | UP000004994 | 34,655 |
| *Populus trichocarpa* | 3694 | UP000006729 | 42,532 |
| *Glycine max* | 3847 | UP000008827 | 55,855 |
| *Brassica rapa subsp. pekinensis* | 51351 | UP000011750 | 40,801 |
| *Arabidopsis thaliana* | 3702 | UP000006548 | 27,476 |

**Table S2.** Cysteine content per plant species. Number of total amino acid residues and cysteine residues for the plant species UniProtKB reference proteomes (see Table S1).

| Species | Total residues | Cysteine residues (%) |
| --- | --- | --- |
| *Ostreococcus tauri* | 3,561,433 | 57,419 (1.61%) |
| *Chlamydomonas reinhardtii* | 13,015,830 | 172,096 (1.32%) |
| *Chara braunii* | 22,684,904 | 361,906 (1.60%) |
| *Marchantia polymorpha* | 7,753,135 | 137,091 (1.77%) |
| *Physcomitrium patens* | 11,734,425 | 215,926 (1.84%) |
| *Selaginella moellendorffii* | 13,317,936 | 253,762 (1.91%) |
| *Amborella trichopoda* | 8,595,683 | 163,976 (1.91%) |
| *Oryza sativa subsp. japonica* | 13,398,719 | 260,517 (1.94%) |
| *Zea mays* | 14,534,640 | 273,307 (1.88%) |
| *Zostera marina* | 7,946,007 | 148,617 (1.87%) |
| *Solanum lycopersicum* | 12,261,183 | 234,815 (1.92%) |
| *Populus trichocarpa* | 15,750,084 | 303,421 (1.93%) |
| *Glycine max* | 21,729,948 | 416,821 (1.92%) |
| *Brassica rapa subsp. pekinensis* | 15,958,951 | 285,707 (1.79%) |
| *Arabidopsis thaliana* | 11,124,147 | 207,864 (1.87%) |

## SUPPLEMENTAL FIGURES

**Figure S1.**
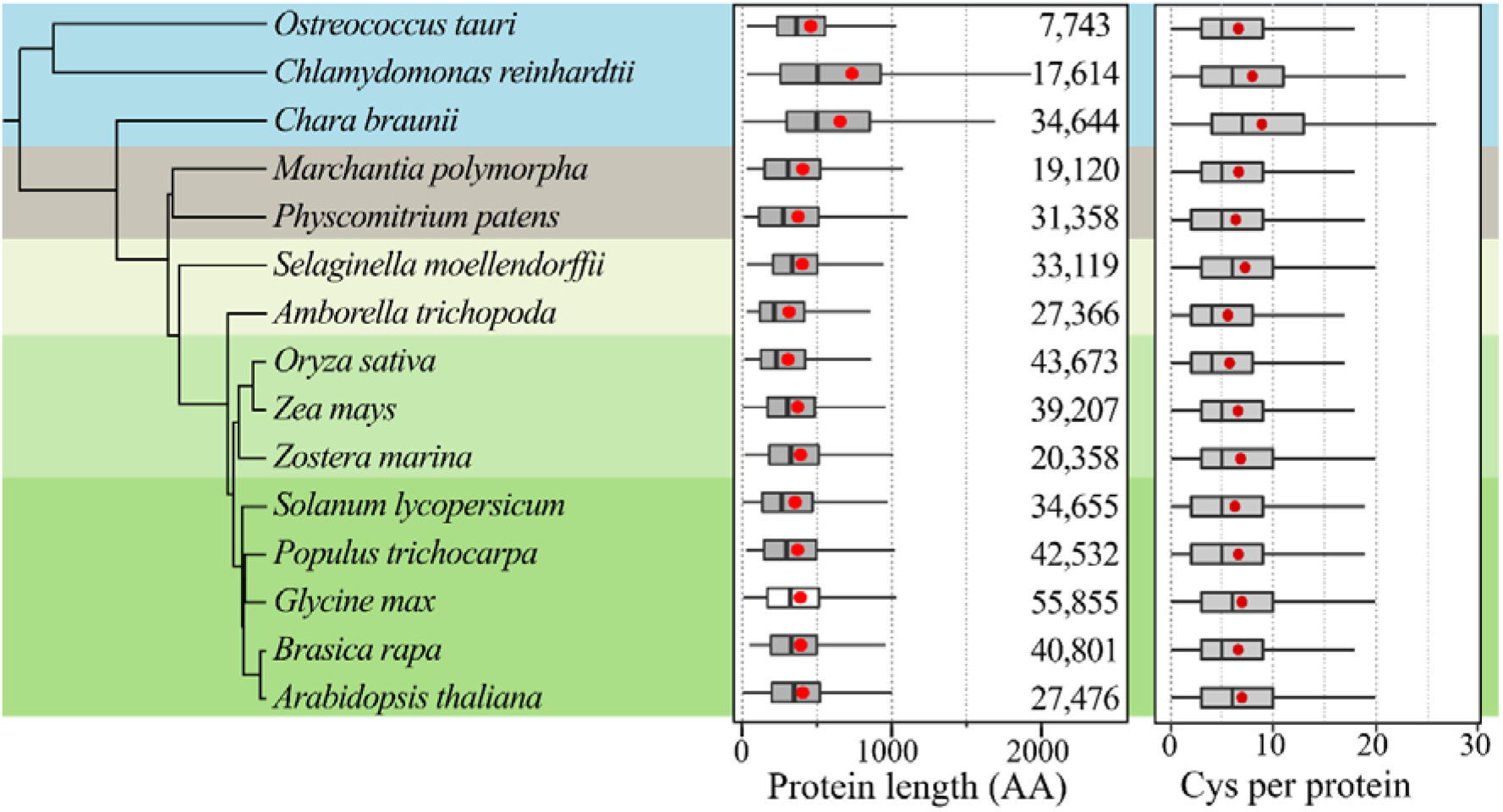
Length and cysteine content per plant species UniProtKB proteome. (*Left*) Schematic representation indicating species relationships. (*Middle*) Protein length distribution per species and total number of proteins. (*Right*) Number of cysteines per protein. Red dots indicate the mean.

**Figure S2.**
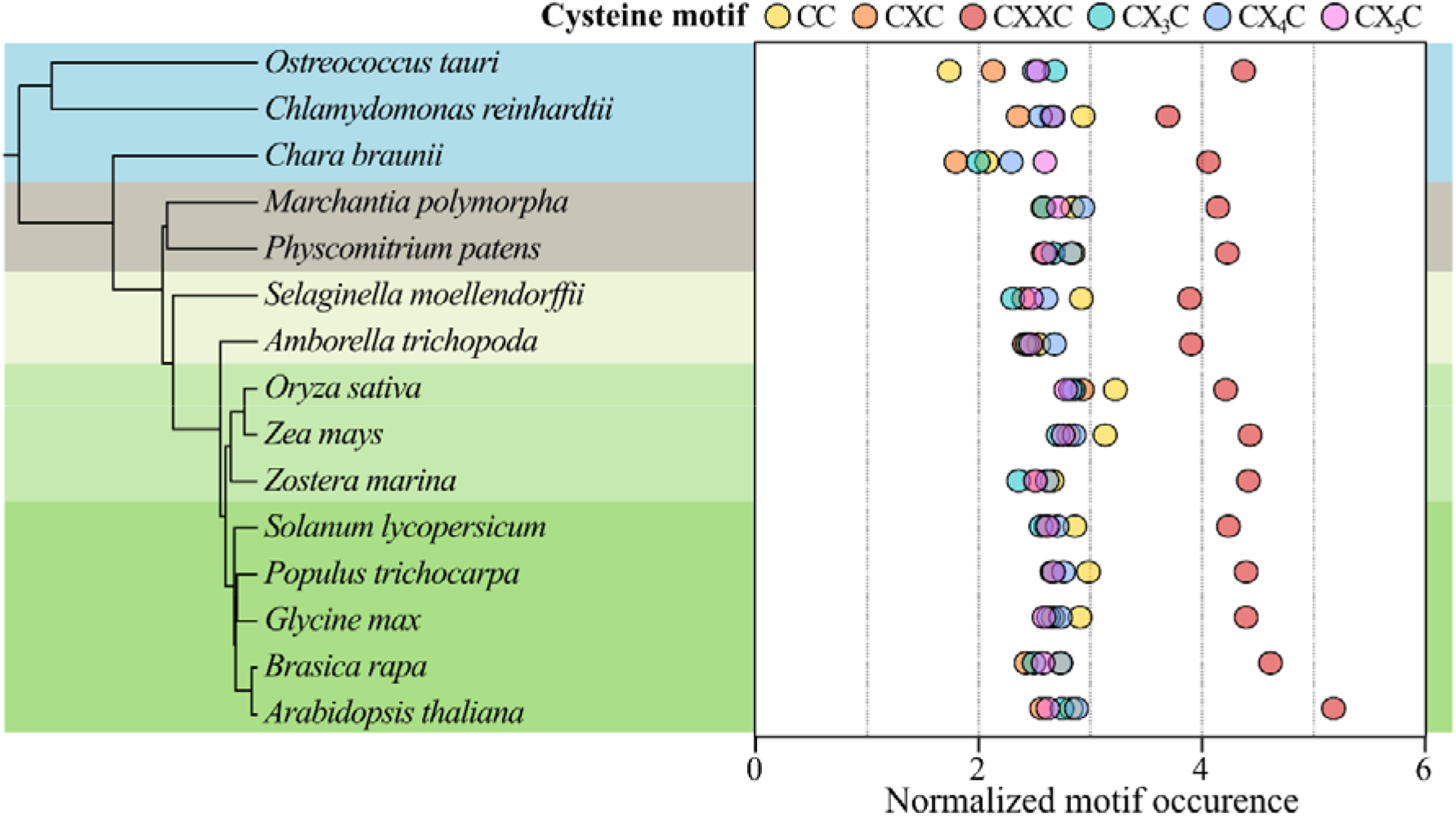
CXXC motifs are overrepresented in plant proteomes. (*Left*) Schematic representation indicating species relationships. (*Right*) Normalized occurrence of CX_n_C motif (n = 0 to 5). The number of CX_n_C motifs was divided by the number of cysteine residues in the proteome and multiplied by a hundred.

**Figure S3.**
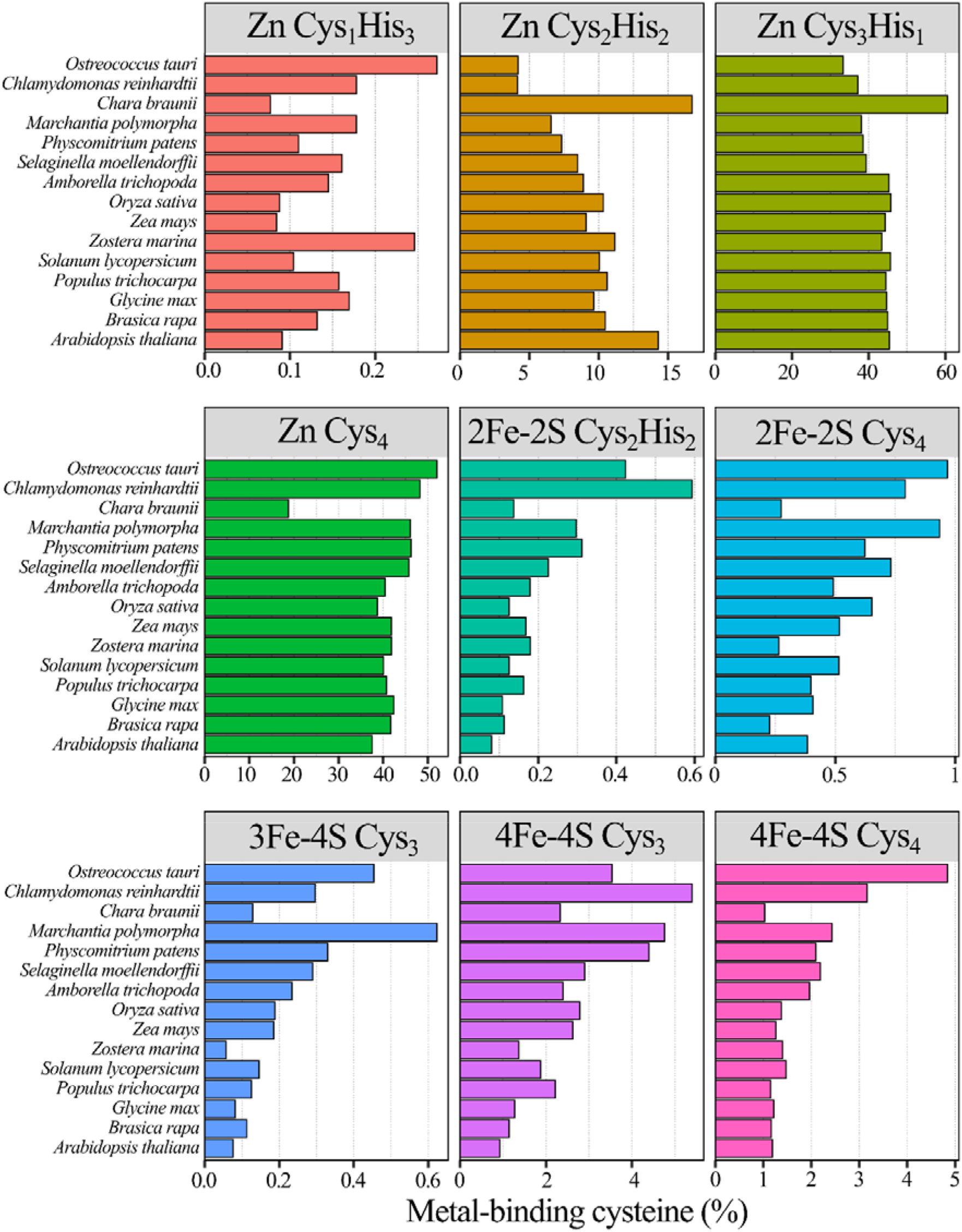
Proportion of cysteines involved in coordination of specific metal ligands. Percentage of cysteine residues participating metal ligand coordination based on AlphaFold2 predictions using the metal ligand search algorithm [40].

**Figure S4.**
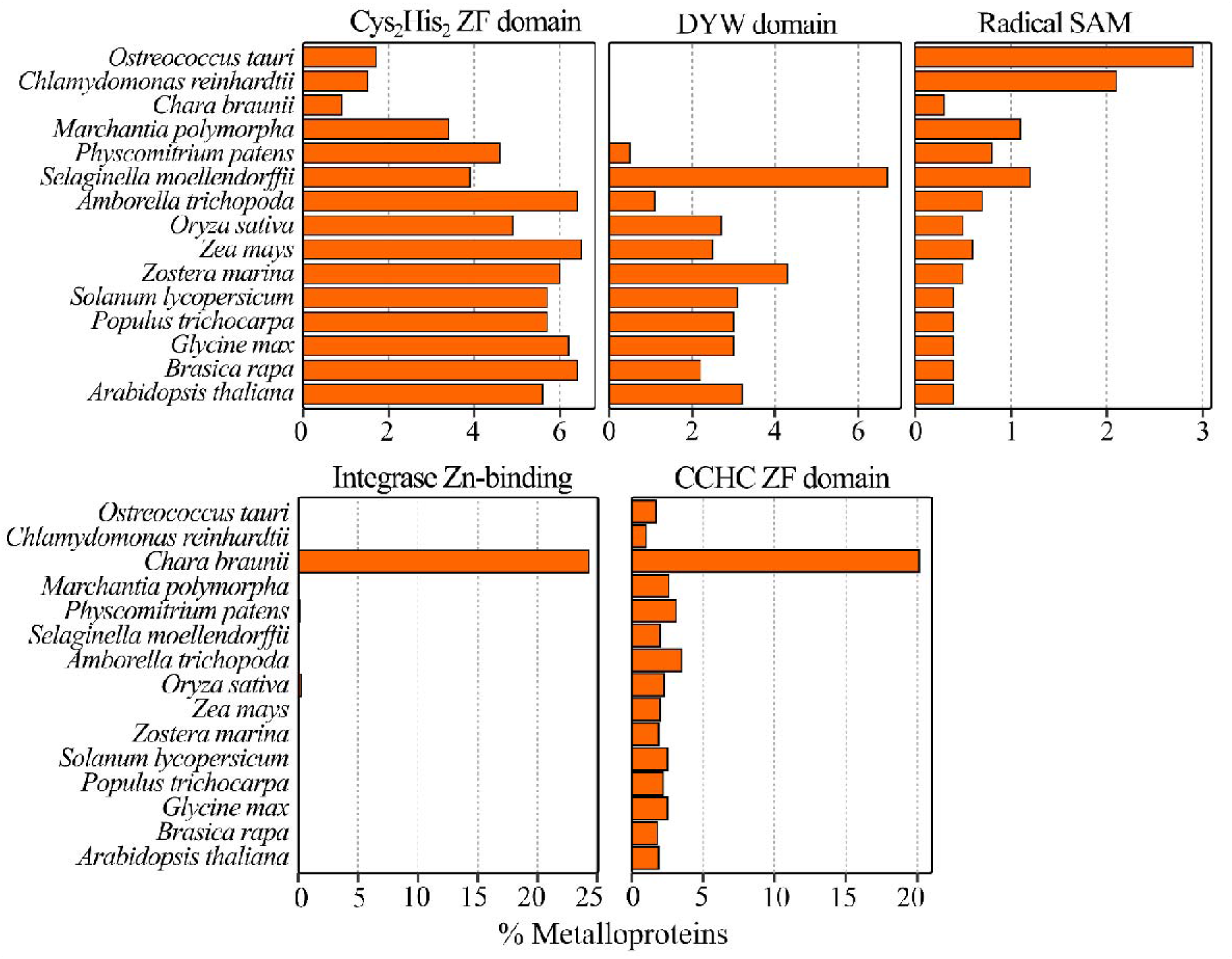
Presence of metal binding InterPro protein domains in AlphaFold2 predicted metalloproteins in the plant kingdom. Proportion of metalloproteins per plant species containing a Cys_2_His_2_ zinc finger domain (IPR013087), DYW domain (IPR032867), radical SAM domain (IPR007197), integrase zinc-binding domain (InterPro IPR041588) or CCHC zinc finger domain (IPR001878). InterPro protein domains [30] are annotated for UniProtKB proteins in all plant species.

**Figure S5.**
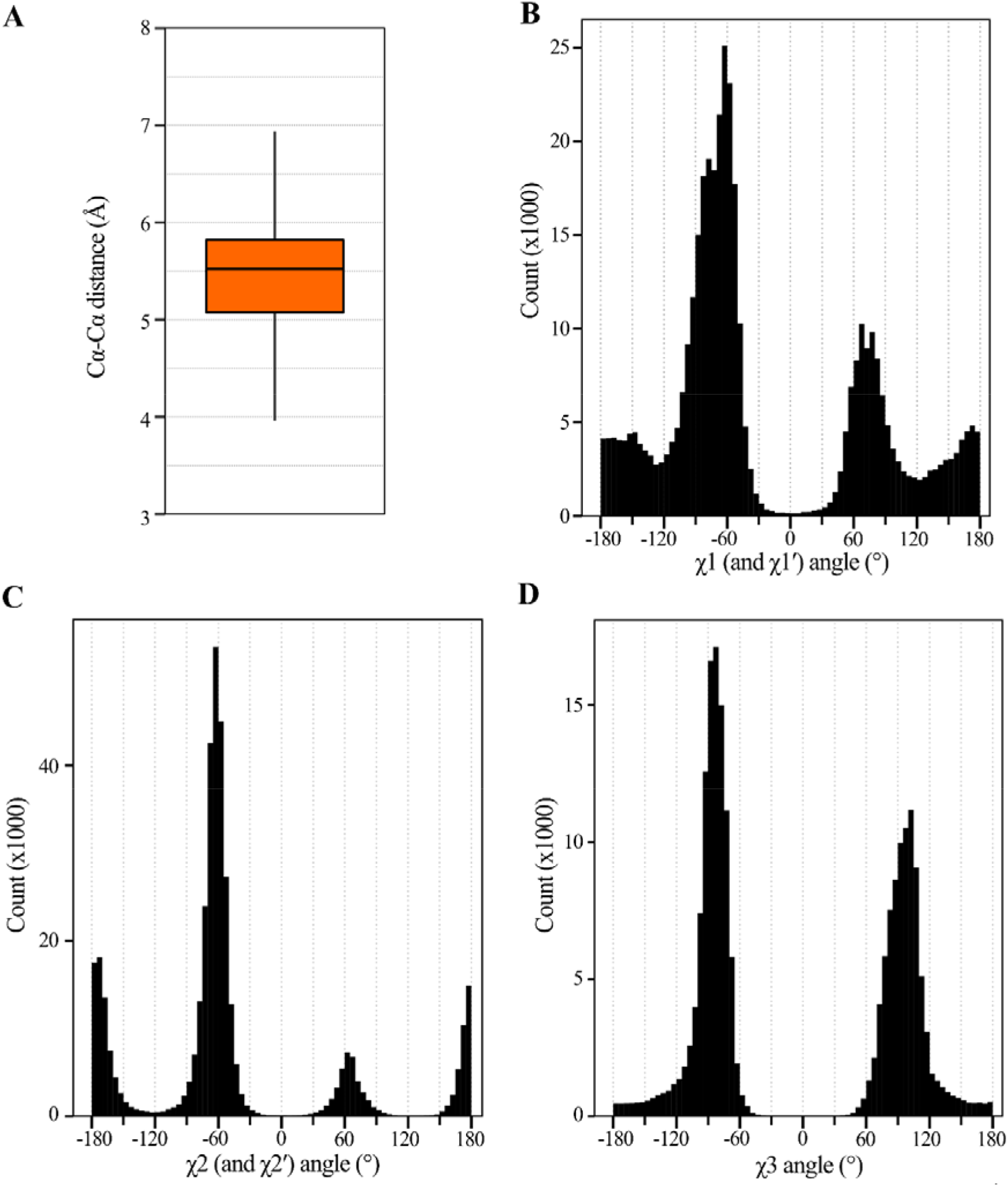
Additional disulfide geometric properties. (**A**) Distribution of cystine Cα-Cα length (Å). (**B-D**) Histograms of χ1, χ2, χ3, χ2′, and χ1′ dihedral angles (x-axis) of disulfides. Angles were binned per 5°.

**Figure S6.**
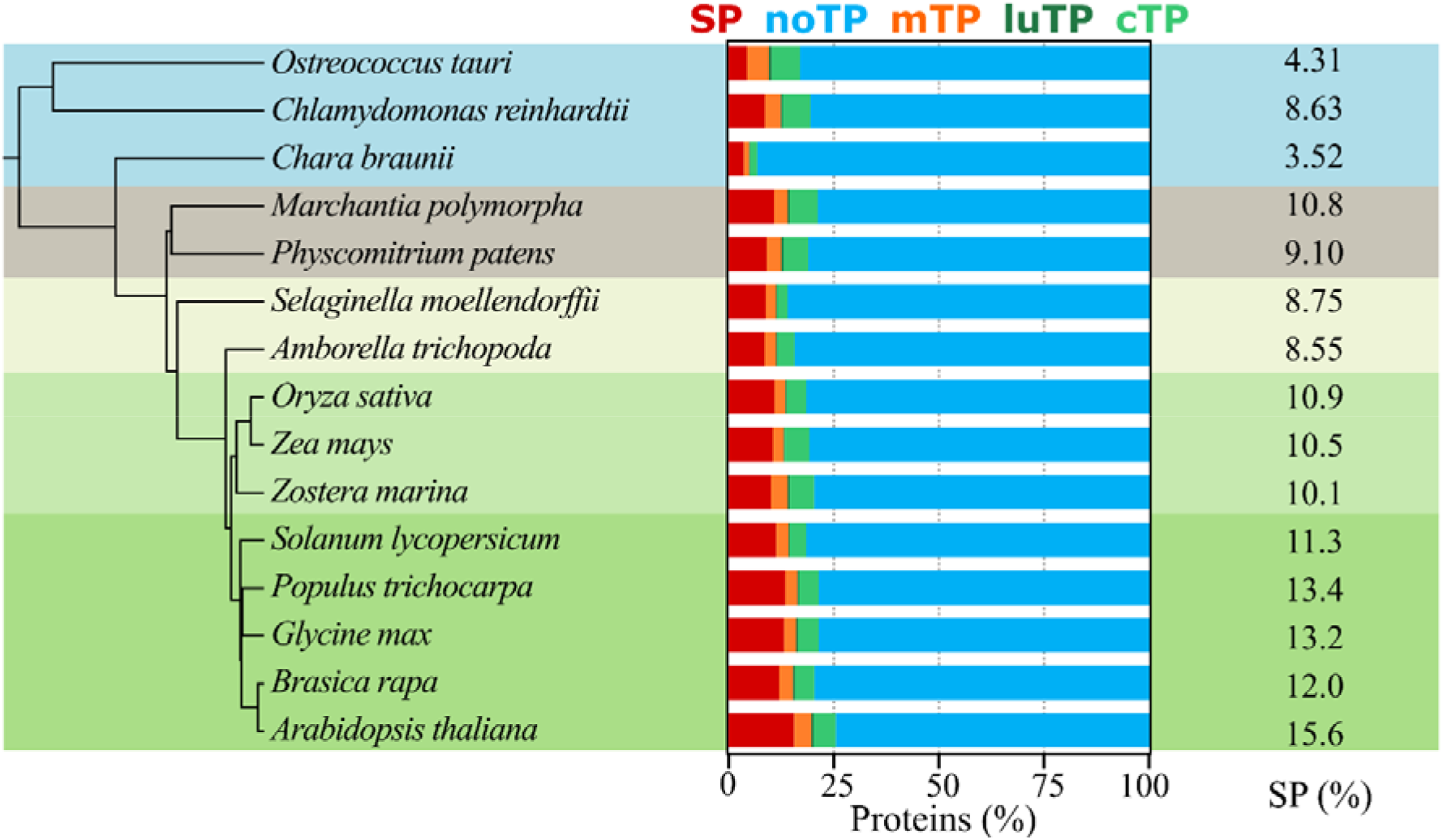
Predicted N-terminal sorting signals per plant proteome. (*Left*) Schematic representation indicating species relationships. (*Middle*) Proportion of proteins containing different targeting peptides as predicted by TargetP 2.0 [31]. (*Right*) Percentage of proteins containing a signal peptide. Abbreviations: cTP, chloroplast transit peptide; luTP, thylakoid lumenal transit peptide; mTP, mitochondrial transit peptide; noTP, no transit peptide; SP, secretory signal peptide.

**Figure S7.**
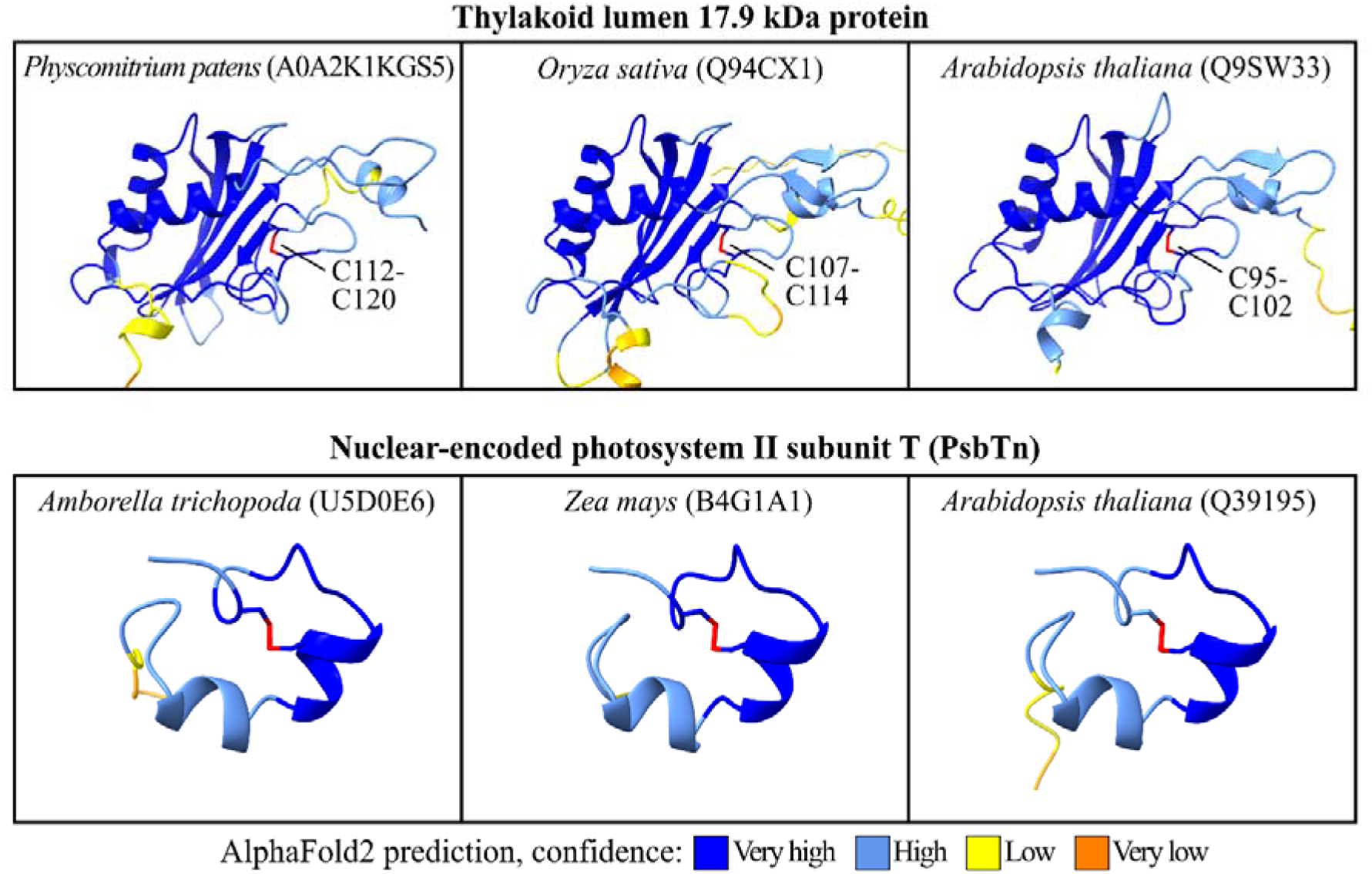
Disulfides in plant thylakoid lumen proteins. AlphaFold2 predicted protein structures with of thylakoid lumen 17.9 kDa protein (*Top*) and the nuclear-encoded photosystem II subunit T (*Bottom*) in different plant species. Disulfides were colored in red. AlphaFold2 predicted proteins were colored by each residue’s pLDDT score, ranging from dark blue (pLDDT > 90, “very high confidence”), light blue (pLDDT > 70, “confident”) and low and orange indicting a “low and very confidence” prediction, respectively. TargetP predicted N-terminal sorting signals were deleted from the structural model.

**Figure S8.**
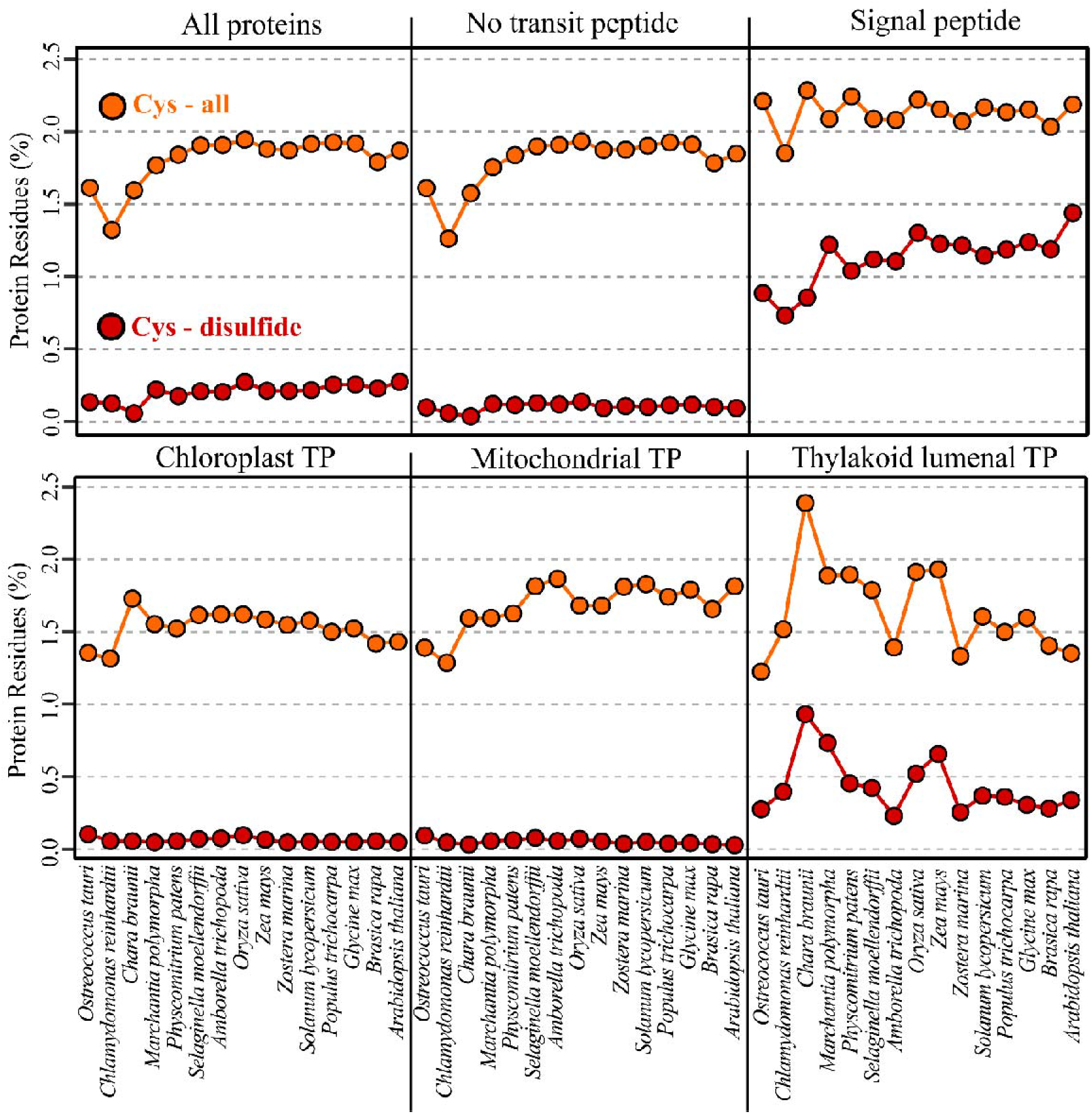
Cysteine content and disulfide formation within subcellular compartments. Proportion of cysteine residues (orange) and cysteine residues forming a disulfide in AlphaFold2 predicted structures (red) in plant proteins according N-terminal sorting signal predictions [31]. Abbreviations: TP, transit peptide.

**Figure S9.**
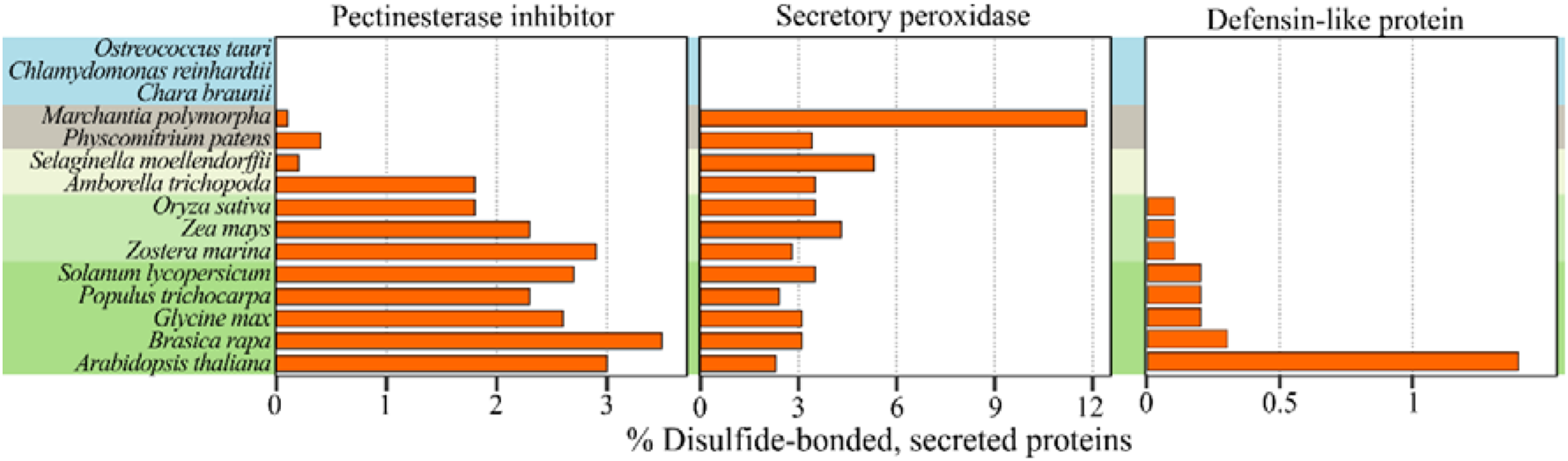
Proportion of secreted proteins (with TargetP 2.0 predicted signal peptide) with predicted disulfide bonds per plant species containing a pectinesterase inhibitor domain (InterPro IPR006501), secretory peroxidase domain (InterPro IPR033905) or belong to the plant-defensin like protein family (InterPro IPR010851).

**Figure S10.**
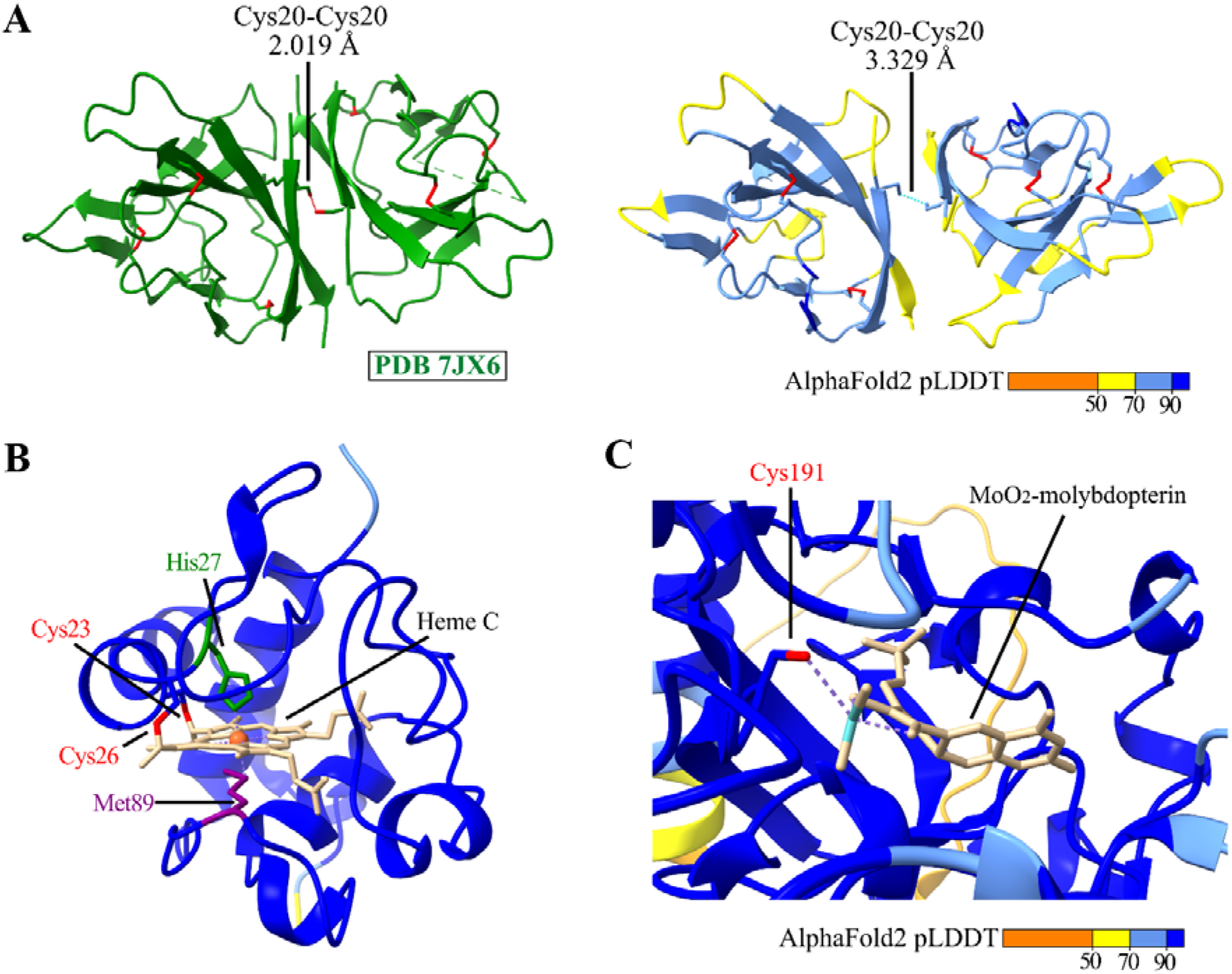
Structural prediction of intermolecular disulfides and cysteine bound cofactors. (**A**) Experimental structure (PDB 7JX6; [74]) and AlphaFold2 predicted model of the SARS-CoV-2 ORF8 homodimer. Disulfides were colored in red. (**B**) AlphaFold2 predicted model of CYTOCHROME C-1 (UniProtKB O23138) with a heme c-type cofactor placed by AlphaFill [75]. (**C**) AlphaFold2 predicted model of NITRATE REDUCTASE2 (UniProtKB P11035) with a MoO_2_-molybdopterin cofactor placed by AlphaFill [75]. AlphaFold2 predicted proteins were colored by each residue’s pLDDT score, ranging from dark blue (pLDDT > 90, “very high confidence”), light blue (pLDDT > 70, “confident”) and low and orange indicting a “low and very confidence” prediction, respectively. TargetP 2.0 predicted N-terminal sorting signals were deleted from the structural model.

